# Elucidating the correlation between the number of TTTTGAT heptamer repeats and cholera toxin promoter activity in *Vibrio cholerae* O1 pandemic strains

**DOI:** 10.1101/2021.05.14.444153

**Authors:** Arindam Naha, Jeffrey H. Withey, Piyali Mukherjee, Rudra Narayan Saha, Prosenjit Samanta, Amit Ghosh, Shin-ichi Miyoshi, Shanta Dutta, Asish K. Mukhopadhyay

**Author notes:** **Corresponding Author: Dr. Asish K. Mukhopadhyay**, Division of Bacteriology, ICMR-National Institute of Cholera and Enteric Diseases, P 33, CIT Road, Scheme XM, Beliaghata, Kolkata 700010, India. Center for Infectious Disease Research, Department of Microbiology and Cell Biology, Indian Institute of Science, Bangalore 560012, India (Piyali Mukherjee).

## Abstract

A complex regulatory cascade controls expression of the cholera toxin genes (*ctxAB*) in *Vibrio cholerae*; which eventually leads to choleragen (CT) production and secretion, resulting in rice watery diarrhoea. The cholera toxin promoter (P*ctxAB*) contains a series of heptad repeats (5’-TTTTGAT-3’); which have been previously shown to play crucial role in *ctxAB* transcriptional regulation by recruiting the transcriptional activators ToxT, ToxR, and the nucleoid-associated protein H-NS along the *ctx* promoter. The numbers of these repeats vary between the two biotypes of *V. cholerae* O1 strains, and even among strains of the same biotype. In this study, we examined P*ctxAB* activation of *V. cholerae* O1 pandemic strains to understand the significance of the distal heptad repeats in regulating *ctx* expression. Interestingly, we found that *ctx* activation may depend on the number of TTTTGAT heptad repeats within P*ctxAB*, and we posit that the occupation of the distal repeats by H-NS could further prevent transcriptional activation of *ctx* genes in *V. cholerae*. We hypothesize that ToxT-dependent transcriptional activation may not require entire displacement of H-NS and propose a revision in the currently accepted model of ToxT dependent P*ctxAB* transcriptional activation.

**IMPORTANCE:** CT production by pathogenic *V. cholerae* O1 strains is regulated through the transcriptional silencing of CTX promoter by H-NS and counter repression by ToxT. The highly AT rich P*ctxAB* is composed of the tandem repeats 5□ TTTTGAT 3□; the numbers of which differ among the classical and El Tor biotypes. However, it is still not very clear whether the numbers of these repeats could be correlated with promoter activation of the *ctx* operon. Here we report the role of the distal repeats in *ctxAB* expression levels. We demonstrate that *PctxAB* activation changes with the number of the heptad repats within the core promoter element, thereby suggesting a model for *ctx* regulation in the toxigenic strains of *V. cholerae*.

## INTRODUCTION

Cholera remains a major health concern in many parts of the world where people lack access to proper sanitation and safe hygiene. Toxigenic strains of the Gram-negative organism *Vibrio cholerae* are the causative agents of this dreadful diarrhoeal disease. Toxin-coregulated pilus (TCP) (1) and cholera toxin (CT) (2) are the major contributing factors to *V. cholerae* colonization and pathogenicity. Expression of *tcpA* and *ctxAB* are controlled by a coordinated network of transcriptional regulators AphA/B, ToxR/S, TcpP/H, ToxT, H-NS (3), and the integration host factor IHF (4). The master virulence activator ToxT is a member of the large AraC/XylS protein family (5). Once produced, ToxT directly activates the transcription of the *ctx* and *tcpA-F* operons (6, 7).

Every ToxT regulated promoter contains one or more copies of a 13-bp DNA sequence called the toxbox; to which ToxT binds and subsequently activates transcription of the downstream genes (8). 5’ portions of the toxboxes are highly conserved and contain a poly (T) tract, whereas the 3’ ends are generally A/T rich. Previously, locations of the toxboxes were identified in the P*ctxAB*, and ToxT binding sites that control *ctxAB* expression were further mapped (9). Despite the presence of several other potential ToxT binding sites within P*ctxAB*, ToxT is recruited to only two toxboxes. P*ctxAB* is composed of highly A/T rich sequences, and contains a series of heptad repeats (5’ TTTTGAT 3’). Interestingly, these repeats are also involved in the binding of global repressor protein H-NS to the P*ctxAB*, which silences *ctxAB* expression under virulence repressing conditions (10). The numbers of these repeats vary among *V. cholerae* O1 strains. Classical biotype strains typically have 6-8 perfect direct repeats, whereas *V. cholerae* El Tor strains most commonly have 4, and at least three direct repeats are essential for ToxT-dependent P*ctxAB* activation (11).

Given that the two functional toxboxes in P*ctxAB* encompass the heptad repeats nearest to the transcriptional start site (TSS), here we set out to determine the significance of the distal TTTTGAT heptad repeats. For this, we have focused on the P*ctxAB* regions of *V. cholerae* O1 pandemic strains with different numbers of heptad repeats to understand their role in transcriptional activation of the *ctxAB* operon. By using *in vitro* CT measurement assays under virulence inducing and repressing conditions (12), P*ctxAB::lacZ* transcriptional fusions, and gel mobility shift experiments, we have also attempted to find out the contribution of the distal repeats in ToxT dependent activation and H-NS mediated transcriptional silencing of the *ctx* regulatory region.

## MATERIALS AND METHODS

### Bacterial strains, plasmids, growth conditions, and antibiotics

All of the strains, plasmids, and oligonucleotides used in this study are listed in Table 1, and were maintained at −80° C in lysogeny broth (LB) containing 25% (v/v) glycerol. All of the *E. coli* strains were grown in LB medium under shaking condition. *V. cholerae* strains were grown overnight in LB medium at 37° C and then subcultured (using a 1:100 dilution) in AKI medium (13) supplemented with or without freshly prepared 0.3% sodium bicarbonate (Na2CO3). P*ctxAB::lacZ* transcriptional fusions for promoter activity assays were constructed by cloning PCR-amplified P*ctxAB* regions form six *V. cholerae* O1 strains in to HindIII-XbaI restriction sites on the reporter plasmid pTL61T (14). For purification of H-NS, 414-bp *hns* gene was PCR amplified from O395 chromosomal DNA by using the primers HNS1 and HNS2 (Table 1). The PCR product was cloned into NdeI-SapI site of the expression vector pTXB-1 (Table 1, New England Biolabs) with a C-terminally tagged chitin binding domain (CBD) to generate pTXB1-H-NS. Following sequencing analysis, recombinant plasmids were introduced into *E. coli V. cholerae* cells. *E. coli* strains (Table 1) were made transformation competent by TSS method (15). *V. cholerae* strains were transformed by electroporation as previously described (16) by using Bio-Rad Gene Pulser (25-μF capacitance, 2.5 kV, and 200 Ω). Following transformation, bacteria were allowed to recover in pre-warmed LB, and plated on LB agar plates containing appropriate antibiotics. Antibiotics were used at the following concentrations: 100 μg/ml ampicillin (Amp), and 25 μg/ml chloramphenicol (Cm).

**Table 1:**
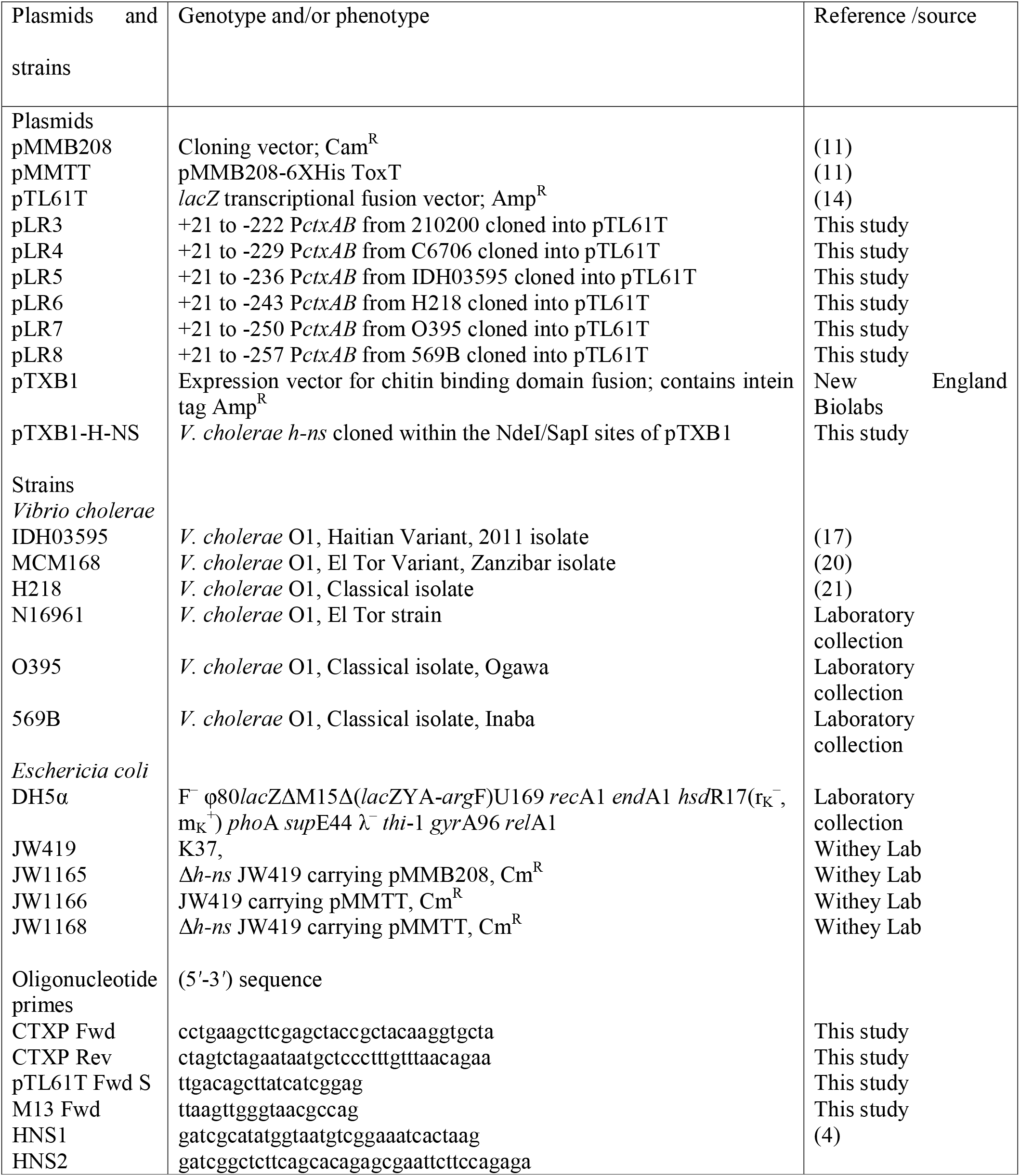
Bacterial plasmids, strains and primer sequences used in the study

### *In vitro* CT measurement and immunoblot analysis

Six *V. cholerae* O1 strains were included in the *in-vitro* toxin measurement study (Table 1). For measurement of CT production by *V. cholerae* O1 strains under *in vitro* conditions, bacteria were cultured in AKI medium under static condition in presence (+) or the absence (-) of bicarbonate (HCO_3_^-^) for 6 hours. The optical densities of the cultures were recorded at 600 nm (O.D. 600nm), and GM1 ganglioside enzyme-linked immunosorbent assays (GM_1_ CT ELISA) were performed as described before (17). A standard curve was generated for every experiment, and CT produced per ml of culture per O.D.600 unit (ng/ml/O.D.600nm) was determined using GraphPad PRISM (V.6.0).

### Promoter activity assay

*ctxAB* promoter activities were determined by measuring the production of ß- Galactosidase level from P*ctxAB::lacZ* transcriptional fusions as previously described (18).

### Protein purification

*V. cholerae* H-NS protein was purified as previously described (4) using the IMPACT-kit (New England Biolabs, Catalogue No. #E6901S). Briefly, protein expression was induced with 0.5 mM isopropyl-β-D-thiogalactopyranoside (IPTG), and bacterial cells were allowed to grow overnight at 16° C. The highly expressed recombinant protein was purified from the cell free soluble fraction of the expression host by using a chitin affinity column. Following on-column cleavage of the CBD by using dithiothreitol (DTT), H-NS was eluted off the column, and the eluted fractions were pooled and subjected to dialysis for 48 hours in 20 mM Tris (pH 8.0), 1 mM EDTA (pH 8.0), 5 mM potassium glutamate, and 0.1 mM DTT. The purity of the H-NS protein was further evaluated through a 15% SDS-PAGE. Concentration of the H-NS protein was measured using Bradford assay (BioRad, USA), and purified protein was stored in aliquots at −80° C in 10% glycerol.

### Electrophoretic mobility shift assays (EMSA)

DNA protein interaction was studied with purified H-NS and DIG labeled PCR amplified P*ctxAB* fragments by using the DIG gel shift kit (2^nd^ Gen, Roche Aplied Science, Mannheim, Germany) as previously described (10). Briefly, DNA fragments C5 (+21 to - 236), and C6 (+21 to −243) were PCR-amplified from P*ctxAB::lacZ* fusion plasmids) with the oligonucleotide pairs CTXP Fwd/CTXP Rev (Table 1). H-NS gel shift assays were carried out in 20μL reactions containing 1 X EMSA buffer (40 mM HEPES, 10 mM magnesium aspartate, 100 mM potassium glutamate, 0.022% NP-40, 100 μg/ml BSA, and 10% glycerol), 25 ng of digoxigenin (DIG)-labeled C5 or C6 DNA, 1 μg of calf thymus DNA, and 180 nM of purified H-NS protein. After incubation at room temperature for 20 minutes, protein-DNA complexes were separated by 5% native PAGE in 0.5X TBE buffer (45 mM Tris-Cl pH 8.3, 45 mM boric acid, 1 mM EDTA) at 80 V, 4°C, and transferred to nylon membrane. DNA was visualized using alkaline phosphatase (AP)-conjugated anti-DIG antibody (Roche) followed by chemiluminescence detection.

## RESULTS

### CT production by *V. cholerae* O1 strains in presence and absence of bicarbonate

HCO3 has been shown previously to increase the activity of ToxT, and thereafter ToxT dependent CT production in *V. cholerae* (19). Cell free culture supernatants from six *V. cholerae* O1 pandemic strains with different numbers of TTTTGAT heptamer repeats within the P*ctxAB* (Figure 1 A) were used to determine any direct corelation between the number of heptad repeats and their relative abilities to produce CT under *in vitro* static AKI conditions in presence and absence of HCO_3_^-^. Strain MCM168, with three perfect TTTTGAT heptad repeats within P*ctxAB* (20), produced the highest amount of CT (~3765 ng/ml/O.D.600nm) in the presence of HCO_3_^-^ (Figure 1B). This El Tor variant strain; which was isolated from Zanzibar, produced CT 9 times higher than N16961 (*P* value 0.01), and approximately 40 fold in excess of the classical strain H218 (*P* value 0.01), which has 6 perfect heptad repeats in the P*ctxAB* (21), and showed a dramatic defect in CT production (~93 ng/ml/O.D.600nm). The other two classical strains 569B and O395 produced around 12-fold (*P* value 0.0001) and 15 times (*P* value 0.04) more CT than H218, respectively. Furthermore, MCM168 produced 2.6-fold heightened CT over both of the classical strains O395 and 569B (*P* value 0.03). IDH03595, a Haitian variant strain isolated from Kolkata with 5 TTTTGAT perfect heptad repeats in P*ctxAB* region (17), produced CT comparable (1827.3 ng/ml/O.D.600nm) to that secreted by 569B (1406.1 ng/ml/O.D.600nm), 1.5 times higher than O395 (*P* value 0.02), and 19.7 fold larger amounts of CT than H218 (*P* value 0.001). IDH03595 demonstrated a more than 4-fold increase in the amount of CT produced in comparison with N16961 (415.3 ng/ml/O.D.600nm, *P* value 0.002), whereas O395 and 569B secreted 2.7 times and 1.5-fold higher CT than N16961 (*P* value 0.0001 and 0.02, respectively).

**Figure 1.**
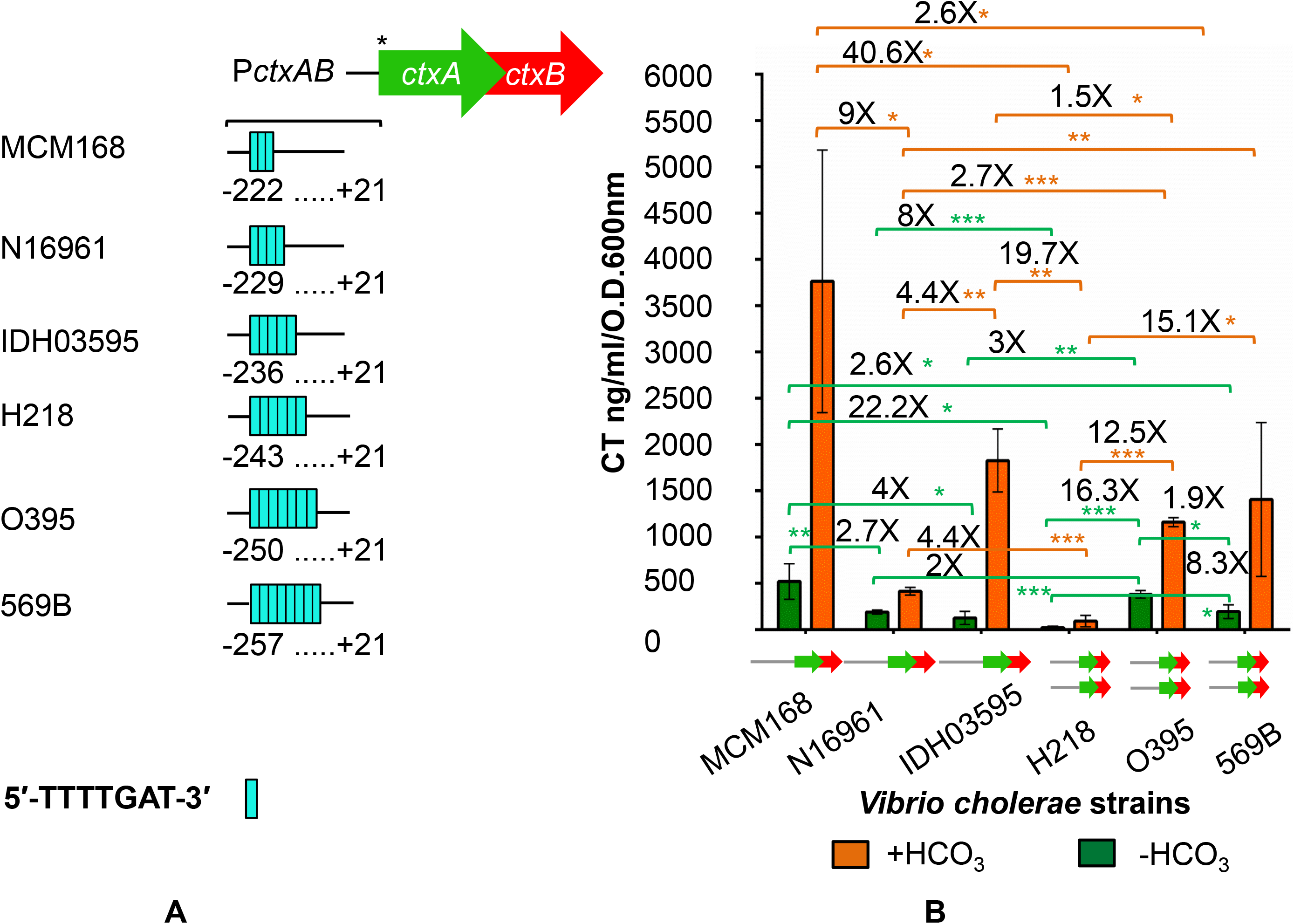
Study of toxin secretion in *V. cholerae*. (**A**) *PctxAB* regions (starting from +21 with 5’ end up to −257) of *V. cholerae* pandemic strains used in this study. Every cyan bar indicates a single TTTTGAT sequence. For better understanding; a graphical view (not drawn to scale) of the *ctxAB* operon has been presented with the first base of the *ctxA* ATG start codon starting from the +22 position (*) relative to the transcriptional start site in the *ctxA* gene (+1). (**B**) Quantification of total CT production by *V. cholerae* O1 strains. Production of CT under AKI conditions in presence and absence of HCO_3_^-^ CT was assessed by GM1 CT ELISA and was measured in ng/ml/OD600. The mean value of at least three individual experiments for each strain is shown with error bars signifying standard deviations. An unpaired two-tailed student’s t test was used to analyse the statistical significance of the data. (* *P* value <0.05, \*\**P* value <0.005, \*\*\**P* value <0.0005). “X” indicates the fold difference in CT values between the indicated strains. The copy number of the *ctxAB* operon is graphically represented over the strain IDs along the X axis.

Interestingly, CT production by MCM168 was found to be the highest (521.525 ng/ml/O.D.600nm) even when HCO_3_^-^ was not included in the AKI media (Figure 1B). Under this condition; the Zanzibar strain produced 2.7 times larger amount of toxin compared to that of N16961 (189.6 ng/ml/O.D.600nm, *P* value 0.002), and 4-fold higher amount of CT than the Haitian variant IDH03595 (~127 ng/ml/O.D.600nm, *P* value 0.006). MCM168 secreted nearly 22-fold heightened CT in comparison with H218; which accumulated barely detectable (23.4 ng/ml/O.D.600nm, *P* value 0.04) holotoxin in the cell free supernatant, and greater than 2.5 times than that produced by the classical strain 569B (194.91 ng/ml/O.D.600nm, *P* value 0.03). The difference in CT secretion between N16961 and H218 appeared 8-fold, where the latter produced significantly lesser amount of CT (*P* value 0.0001) than the former one. The classical strain O395 produced 382.7 ng/ml/O.D.600nm of CT, which is approximately 16 times increased compared to H218 (*P* value 0.0001), 3-fold larger than compared with IDH03595 (*P* value 0.001), and twice that produced by N16961 (*P* value 0.0001). Another one of the three classical strains included in CT-ELISA experiments; 569B, secreted more than 8-fold amount of higher CT than H218 (*P* value 0.03), and half of the amount of toxin produced by O395 (*P* value 0.01) in absence of HCO_3_^-^. Given that H-NS is expressed at similar levels in both El Tor and classical strains (22) and the level of intracellular ToxT protein differs significantly among *V. cholerae* isolates (23), we anticipated that at least ToxT independent (-HCO_3_^-^) heightened CT production in MCM168; which has a single copy CTX prophage (20), could be attributed to weaker H-NS mediated repression on P*ctxAB*. However, our data did not allow us to directly correlate ToxT regulated (+HCO_3_^-^) CT production by the six *V. cholerae* strains with the differing occurrences of TTTTGAT repeats upstream of the *ctxAB* operon.

### TTTTGAT repeats participate in regulating P*ctxAB* activity

Previously, the heptad repeat sequences had been shown to be important for H-NS binding within *PctxAB* (10). Our CT ELISA data motivated us to investigate whether the higher number of the heptad repeats in classical strains could result in increased *ctx* transcriptional repression by H-NS, and thereby in reduced promoter activity. Classical and El Tor biotype strains differentially regulate production of CT and TCP (24). To begin characterizing the role of the heptad repeats in *ctx* activation; P*ctxAB::lacZ* transcriptional fusion plasmids (pLR3-pLR8, Table 1) were constructed by placing varying lengths of *ctx* regulatory regions upstream of a promoter-less *lacZ* gene in pTL61T. The fusion constructs contain P*ctxAB* with three to eight TTTTGAT tandem repeats within the intergenic space between the *ctxAB* operon and the *zot* gene. (Figure 2), amplified from the El Tor variant strains MCM168, El Tor strain N16961, Haitian variant isolate IDH03595, classical strains H218, O395, and 569B as described under “Materials and Methods”. Each of these P*ctxAB::lacZ* fusions has a 3’ end extended to +21 (relative to the transcription start site), and contains the −10 and −35 basal promoter element, and two toxboxes which consist of three perfect TTTTGAT promoter proximal repeats and the DNA sequence 5’- GATTTCAAATAAT-3’ (9). The 5’-end-points of these P*ctxAB::lacZ* fusions are: —222, – 229, –236, –243, –250, and –257. As the number of the heptad repeats ranges between 3-5 in El Tor strains and 6-8 in classical biotype strains, P*ctxAB::lacZ* transriptional fusion plasmids pLR3 (+21 to −222), pLR5 (+21 to −236), pLR6 (+21 to −243), and pLR8 (+21 to −257) were tested for their abilities to drive synthesis of a promoterless *lacZ* gene after they were propagated in *V. cholerae* strain N16961. Interestingly, we found that the promoter activities of the fusion plasmids dropped gradually as the number of the heptad repeats increased within the P*ctxAB::lacZ* fusions (Figure 3). P*ctxAB* construct pLR3 (upstream end −222) with 3 TTTTGAT heptad repeats produced the highest level of β-galactosidase activity (1767.7 Miller Units), which was 1.6-fold higher compared to pLR5 (*P* value 0.04) with 5 TTTTGAT heptamer repeats. This observation is in agreement with the previous work (11) that demonstrated P*ctxAB* constructs extending to upstream end-point −76 relative to the transcription start site is sufficient for ToxT dependent activation of the *ctx* genes. The promoter activity further declined in pLR6 (in which the P*ctxAB* insert extends till −243 with 6 perfect heptad repeats) by 1.3 times in comparison with pLR5 (*P* value 0.003)) and more than two fold than that of pLR3 (*P* value 0.001). A drastic loss in *ctx-lacZ* expression could be observed with the P*ctxAB::lacZ* construct pLR8, which extends up to −257 at the 5’ end with 8 consecutive heptad repeats. pLR8 produced more than 2.5-fold less β-galactosidase activity as compared to pLR5 (*P* value 0.0001), and 5.9-fold less than pLR3 (*P* value 0.0005). These results suggest that the degree of P*ctxAB* activation perhaps depends on the number of heptad repeats, and the distal TTTTGAT heptad repeats could play a part in H-NS binding and *ctx* repression.

**Figure 2.**
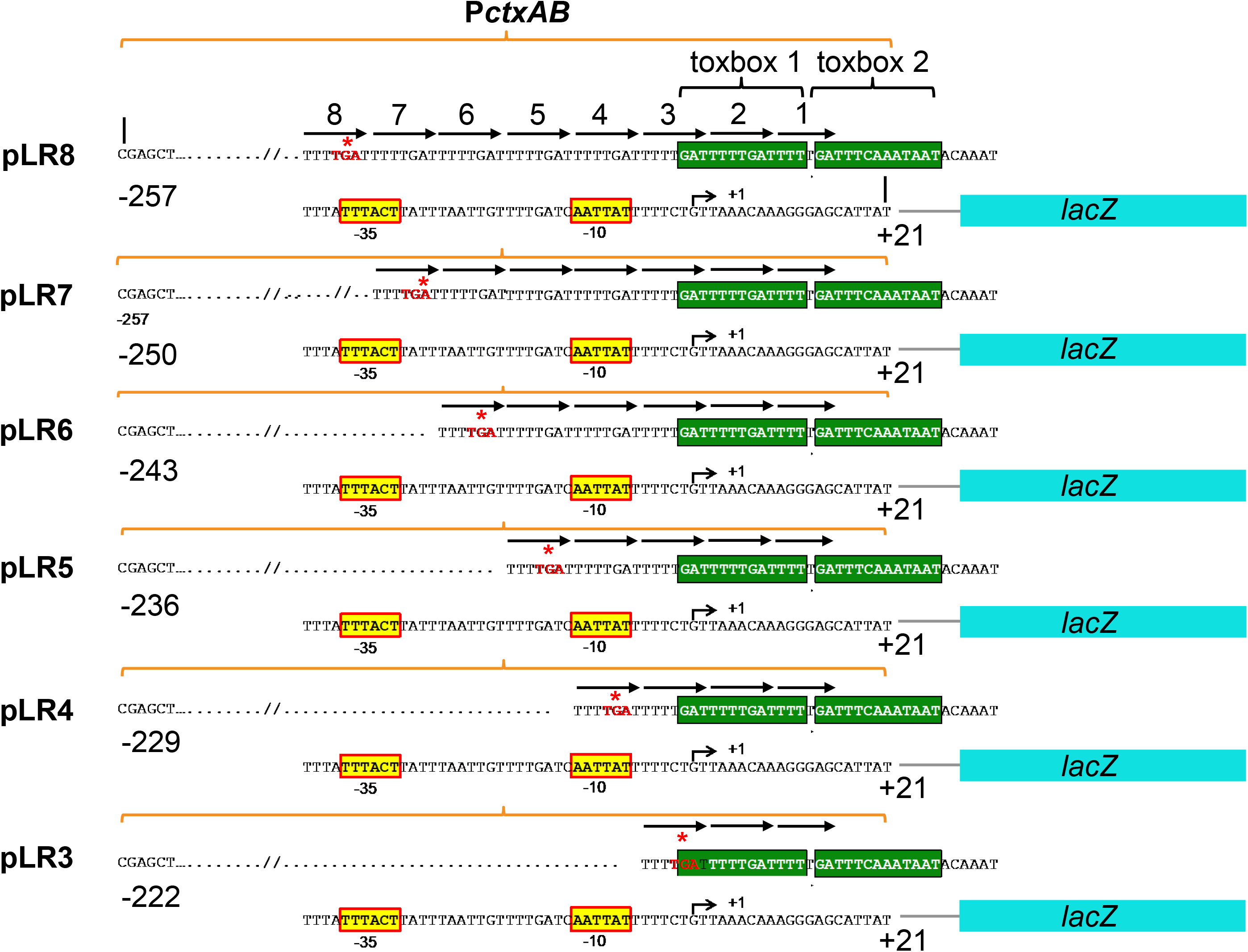
P*ctxAB::lacZ* transcriptional fusion plamids. P*ctxAB::lacZ* transcriptional fusion plasmids containing *Zot-ctxA* intergenic regions (starting from +21 with 5’ end up to −257) amplified from *V. cholerae* pandemic strains. The ToxT binding toxboxes; toxbox 1 and toxbox 2, are indicated by green boxes. The bend arrow illustrates the transcription start site (+1) in the *ctxA* gene. Putative −10 and −35 core elements are highlighted within the yellow boxes. TGA stop codon of the *zot* gene has been marked with the red asterisk. TTTTGAT heptad repeats are marked with black arrows, and have been numbered from the promoter-proximal end. P*ctxAB* inserts are ligated to an otherwise promoterless *lacZ* gene (cyan boxes); the transcription of which is directed from the upstream *ctxAB* regulatory regions following the addition of the inducer in the media as described under “materials and methods”.

**Figure 3.**
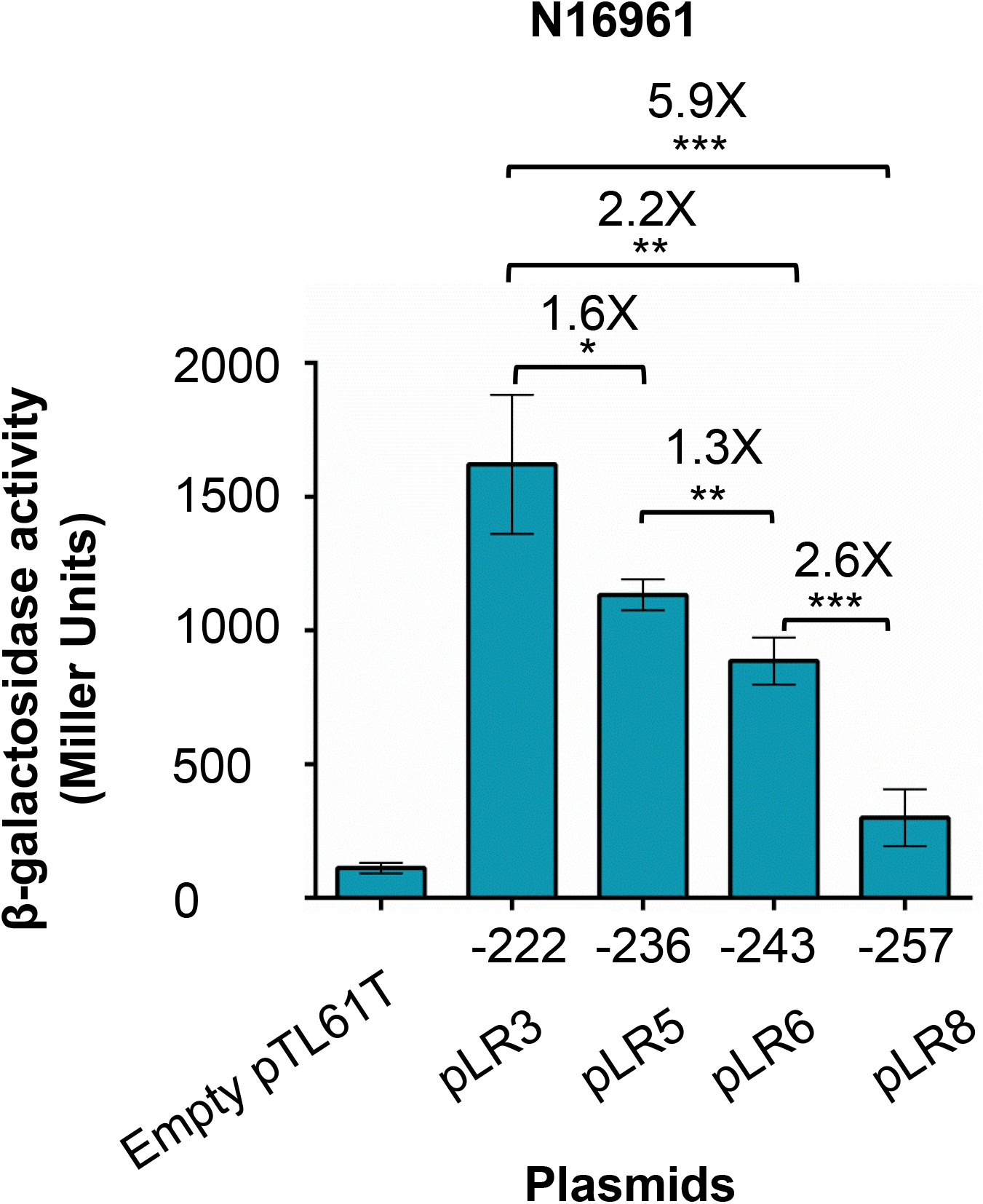
β-galactosidase assay for measuring *PctxAB* activity. Derivatives of WT N16961 (*hns^+^toxT^+^*) were grown to mid-exponential phase in LB medium at 30° C under shaking conditions and tested for their abilities to produce β-galactosidase from the *ctx::lacZ* fusion constructs. Data are expressed in Miller units (MU) and the mean values of at least three individual experiments are plotted. Error bars indicate standard deviation for each group. Asterisks indicate the statistical significance of the data determined by two-tailed student’s t test (* *P* value <0.05, \*\**P* value <0.005, \*\*\**P* value <0.0005). “X” indicates the fold difference MU between the indicated strains. The 5□ ends of the *PctxAB::lacZ* fusions are indicated along the X axis over the respective reporter plasmids.

### DNA binding by H-NS

To understand the importance of the distal heptad repeats within classical P*ctxAB* in H-NS binding, we aimed to study DNA-protein interaction with *ctx* promoter fragments amplified from H218, which has the minimum number of the heptad repeats among the classical strains included in the CT-ELISA experiments and demonstrated severely compromised CT production. We affinity purified full length H-NS protein (15.1 kDa) from *E. coli* using chitin slurry (Supplementary figure 1) as described in “Materials and Methods”. For studying interactions between H-NS and the distal heptad repeats, gel shift assays (Figure 4B) were performed with 180 nM of purified H-NS, and DIG labelled fragments C5 (five TTTTGAT repeats) or C6 (six TTTTGAT repeats) in 1X EMSA buffer. Although both of the DNA fragments interacted strongly with H-NS, we could observe that a supershift appeared when purified H-NS was added to C6, which extends till −243 at the 5’ end. Interestingly, this supershift was not obviously present when the C5 DNA substrate with five TTTTGAT heptad repeats was used as probe. Importantly; unbound free C6 DNA could also be observed in the EMSA blot even in presence of H-NS. The above-mentioned observations suggest that probably the distal heptad repeats 6-8 contribute to stronger H-NS repression than those proximal to the promoter and, therefore, transcriptional silencing of P*ctxAB* may inversely correlate with the number of the distal heptads within the *ctx* regulatory region.

**Figure 4.**
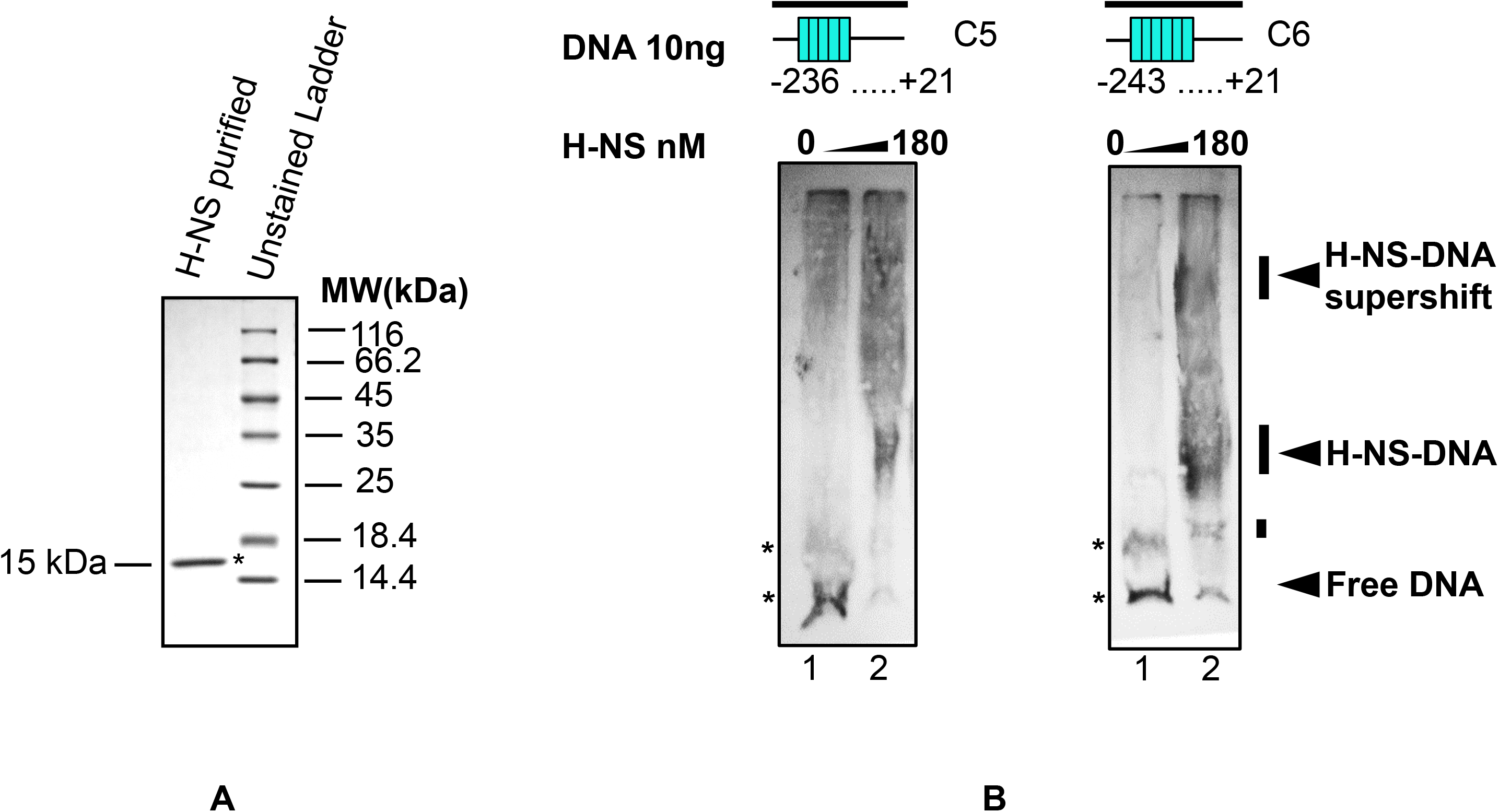
Gel shift assay. (**A**). Purified H-NS protein was used for EMSA experiments. Positions of the marker proteins (Pierce^™^ Unstained Protein MW Marker, Catalogue No. 26610) of known molecular weights (kDa) are indicated. (**B**). Interaction of H-NS with the distal heptad repeats. The DIG-labeled P*ctxAB* fragments C5 (+21 to −236) and C6 (+21 to −243) used in this experiment are schematically illustrated. Each cyan bar represents a single TTTTGAT heptad sequence. 25 ng each of the DIG-labelled fragments were incubated with purified 180 nM H-NS and analysed on a nondenaturing polyacrylamide gel. Lane 1 in each set is devoid H-NS, whereas lane 2 contains 180 nM of H-NS. Blots were probed with HRP conjugated anti-DIG antibody followed by visualization using chemiluminescence.

### Regulation of P*ctxAB* activity by ToxT and H-NS

P*ctxAB* is highly A/T rich (86%) and composed of tandem repeats of the sequence TTTTGAT. The TTTTGAT heptad repeats were previously identified as binding sites for the transcriptional activators ToxR (25), (26), and ToxT (7), (9). For further understanding of the roles of the heptad repeats in *PctxAB* transcriptional activation by ToxT and H-NS, experiments with *Δhns* mutants *V. cholerae* were attempted. However, despite repeated attempts we were unable to obtain a viable *Δhns* mutant. Therefore, we carried out promoter activity experiments in *Δhns E coli* strains with either recombinant plasmid pMMTT expressing 6-His-ToxT upon IPTG induction or the empty vector pMMB208 alone (Table 1, (11). Using three different lots of *E. coli* strains, promoter activities with fusion plasmids pLR3-8 were estimated in the presence and absence of both ToxT and H-NS, thereafter eliminating any interfering effect of ToxR, which also utilizes the heptad repeats to directly bind the P*ctxAB* (26).

Consistent with our previous results, the P*ctxAB::lacZ* reporter plasmid pLR3 produced the highest level of β-galactosidase activity (5818.7 Miller Units) in H-NS^+^ ToxT^+^ *E. coli* cells (Figure 5), resembling the scenario when both H-NS and ToxT (in active form) are present inside *V. cholerae*. While we could found no significant change in promoter activity from pLR4, P*ctxAB::lacZ* expression from pLR5 showed a reduction of 1.7- fold (3338 Miller Units, *P* value 0.02) in β-galactosidase level compared to pLR3, and 1.3 times than that from pLR4 (*P* value 0.01) which produced 4394.4 Miller Units of β-galactosidase activity. The *PctxAB* activity was further decreased by a magnitude of 1.2 (*P* value 0.0001) in the fusion construct pLR6, which produced Miller Units of activity 1.7 times lower than that produced by pLR4 (*P* value 0.0001), and approximately two times less as compared to pLR2 (*P* value 0.002). The other two transcriptional fusion plasmids pLR7 and pLR8 showed near identical levels of promoter activity produced by pLR5 (3053.8 and 2921.3 Miller Units of β- galactosidase activity respectively). Empty pTL61T plasmid; which was used as vector control (mimicking the virulence repressing condition in *V. cholerae* when ToxT activity is not present to relieve H-NS repression), produced ~460 Miller units of β-galactosidase activity.

**Figure 5.**
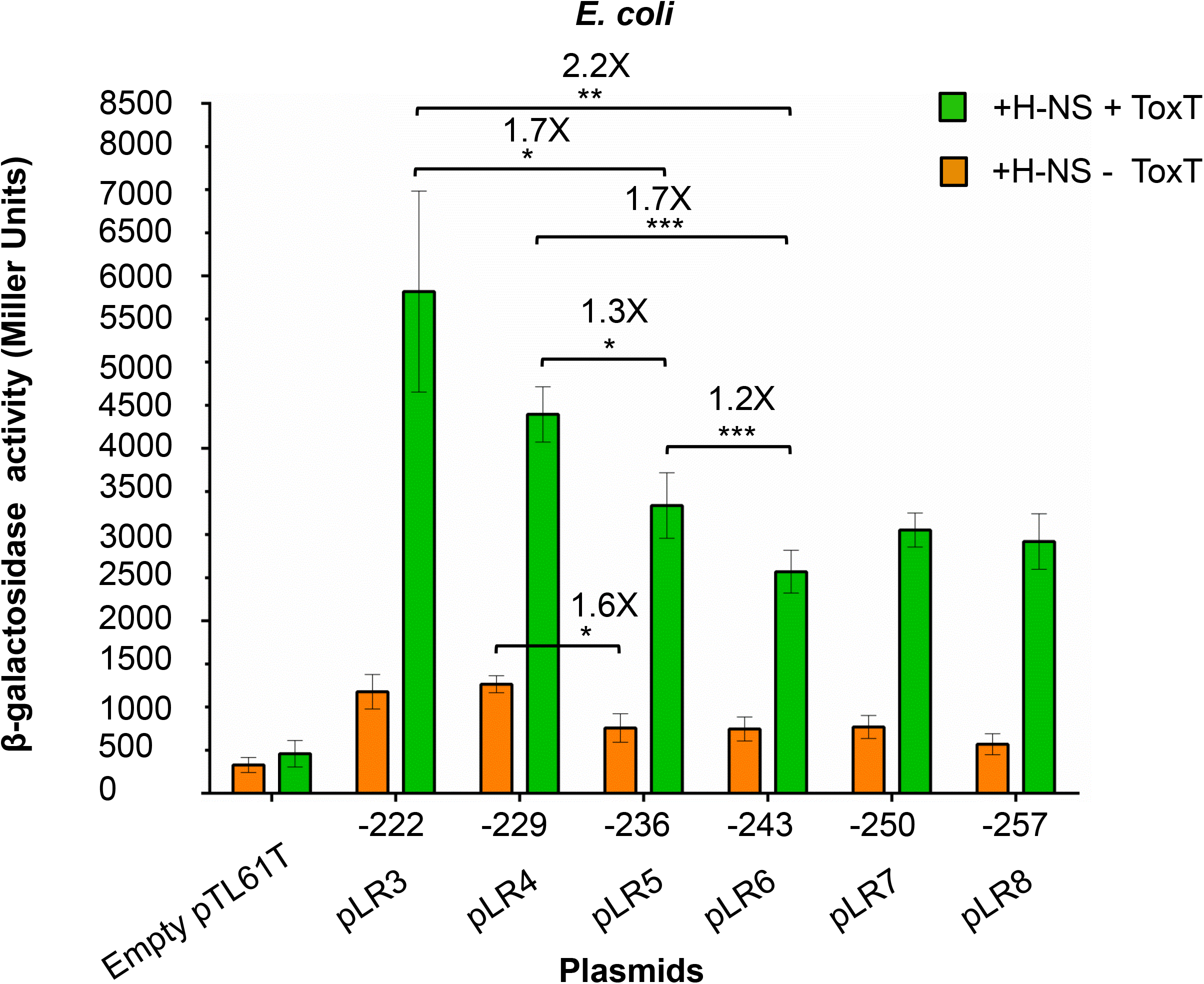
Analysis of *ctx::lacZ* fusions in *E. coli*. Derivatives of *E. coli* K37 strain JW419 transformed with recombinant plasmids carrying P*ctxAB* fusion constructs were grown at 37° C overnight, and diluted 1:100 in fresh LB medium supplemented with 1 mM IPTG and were allowed to grow at 30° C for 3 hours. β-Galactosidase activity was measured in Miller Units (MU) as described previously (17). A minimum of three individual experiments for each strain were performed, and mean values were plotted to with error bars signifying standard deviations for each strain. An unpaired two-tailed student’s t test was used to analyse the statistical significance of the data. (* *P* value <0.05, \*\**P* value <0.005, \*\*\**P* value <0.0005). “X” indicates the fold difference in MU between the indicated strains. +H-NS +ToxT represents *E. coli* strain JW1166 expressing 6XHis ToxT from the IPTG inducible plasmid pMMTT (*toxT* gene cloned into pMMB208 under the control of a *tac* promoter). +H-NS-ToxT refers to *E. coli* strain JW1165 carrying empty plasmid pMMB208. The 5□ ends of the *PctxAB::lacZ* fusions are indicated along the X axis over the respective reporter plasmids.

The result of these experiments indicated that *ctx* activation and the number of the distal repeats possibly correlate. In all of the *E. coli* strains (H-NS^+^), little promoter activity could be observed without ToxT (Figure 5). Interestingly, promoter activity from pLR5 dropped by 1.6-fold (619.3 Miller Units) compared with pLR3 and pLR4 (*P* value <0.05), indicating that this region; which extends up to −236 and consists of five direct TTTTGAT heptad repeats, could be a potential H-NS binding site. However, no further change could be observed with any of the other P*ctxAB::lacZ* transcriptional fusions, thereby indicating that not all of the perfect repeats possibly could recruit H-NS to P*ctxAB* with equivalent affinity. Our results with the *E. coli* strains suggest that the degree of H-NS binding and *ctx* repression, and therefore transcriptional activation from the P*ctxAB* corelate with the number of TTTTGAT heptamer repeats, albeit following a less straight forward pattern.

The above-described fusion plasmids were also mobilized into *E.coli hns* strains expressing 6XHis-ToxT from an IPTG inducible plasmid to determine the role of the heptad repeats in ToxT dependent transcriptional activation. Once ToxT was expressed, P*ctxAB* activity heightened dramatically (Figure 6). Promoter activity was strikingly similar in all of the reporter fusions except pLR3, which produced β-galactosidase activity (~12286 Miller Units) 1.3 times higher than pLR4 and the other constructs (*P* value 0.02). This particular result supports previous findings (9) that ToxT was not observed to bind to the distal heptad repeats in footprinting experiments, and indicated that P*ctxAB* is activated by ToxT even in the *E. coli* background, possibly through stimulating RNA-polymerase (RNAP) when H-NS is absent (11).

**Figure 6.**
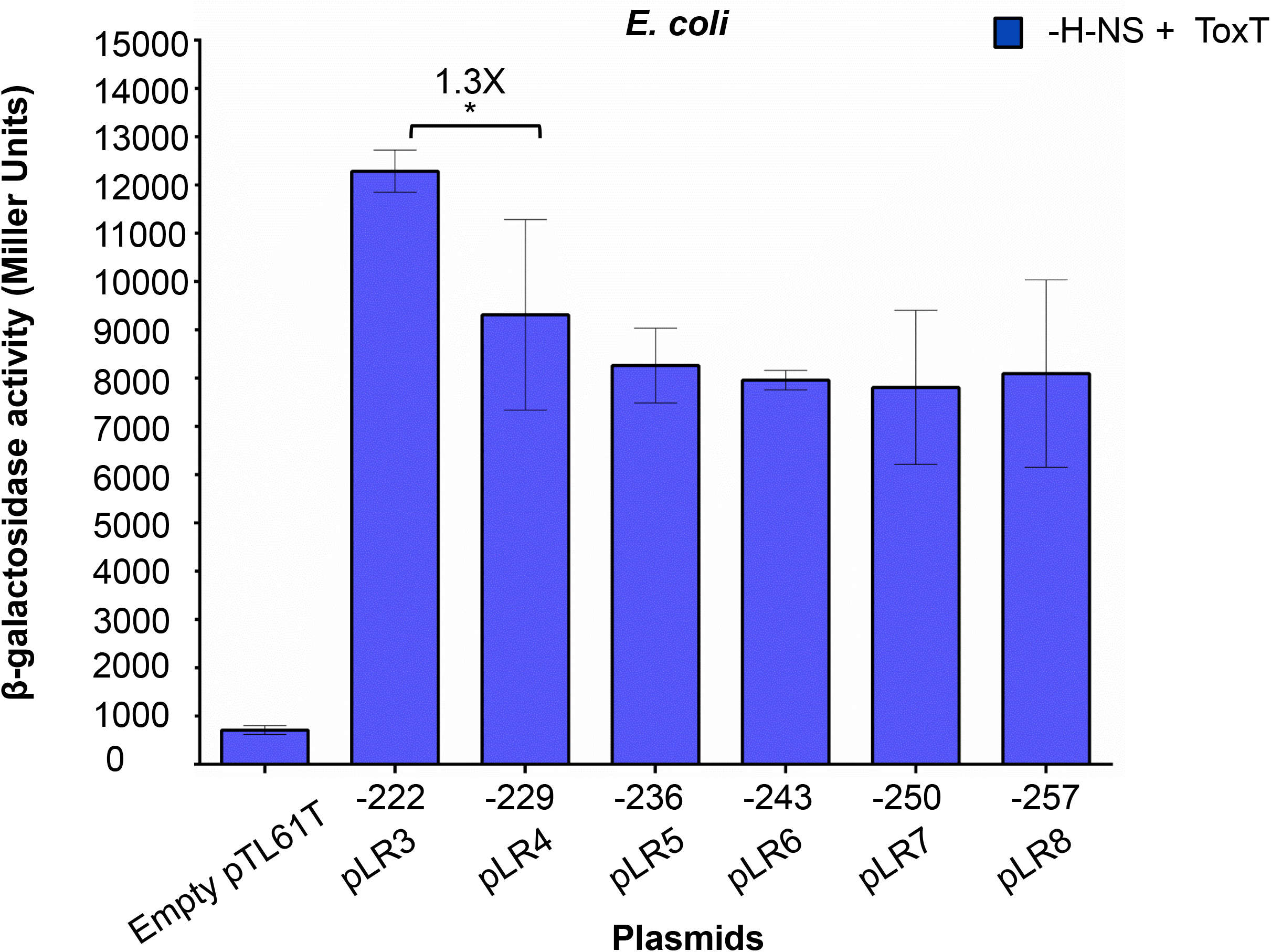
Analysis of ToxT dependent P*ctxAB* in *Δh-ns E. coli*. β-Galactosidase activity assays were performed with indicated *ctx::lacZ* reporter plasmids and lysates of the *E. coli* K37 strain JW1168. Promoter activations were reflected by Miller Units (MU). -H-NS +ToxT represents *h-ns* disruption and ToxT complementation from the pMMTT (*toxT* gene cloned into pMMB208 under the control of a *tac* promoter). +H-NS-ToxT refers to *E. coli* strain JW1165 carrying empty plasmid pMMB208. Error bars represent standard deviations from a minimum of three biological replicates, and asterisk signify the *P* value (* *P* value <0.05). “X” indicates the fold difference in MU□. The 5□ ends of the *PctxAB::lacZ* fusions are indicated along the X axis over the respective reporter plasmids.

Taken together, our results imply that the distal repeats have a role in negatively regulating P*ctxAB* activity, and this observation was additionally supported by our *in vitro* EMSA experiments.

## DISCUSSION

Coordinated expression of *ctxAB* is regulated by a complex pathway in which both specific regulators, such as ToxR/S, TcpP/H, and ToxT, and global regulators, such as H-NS and integration host factor (IHF), influence the expression of P*<u>ctxAB</u>*. (6), (27), (28) (29), (4). Once ToxT is produced, it activates the transcription of the virulence associated genes including the *ctxAB* operon (7).

*ctx* mRNA expression is repressed by the 15-kDa histone-like nucleoid associated protein H-NS, which binds to A/T rich DNA sequences in a process known as ‘xenogeneic silencing (30). H-NS recognizes a 10 bp high-affinity binding motif, TCGATAAATT (30), and eventually polymerizes along the AT-rich DNA sequence by utilizing TTTTGAT heptad repeats within P*ctxAB* (10). The numbers of these heptad repeats, which varies between the classical and El Tor biotypes, have earlier been identified as binding sites for positive regulators ToxR (25) (26) and ToxT (23) (9). The level of ToxT production differs across *V. cholerae* strains (32), as does the amount of ToxT dependent CT production under *in vitro* virulence inducing conditions. In this study, experiments were conducted to elucidate the role of the distal heptad repeats in regulating P*ctxAB* transcriptional activation. Induction of ToxT activity by HCO_3_^-^ globally results in enhanced transcriptional activation of ToxT activated genes, which includes *ctxAB* (19). Therefore, six pandemic strains of *V. cholerae* with different numbers of TTTTGAT heptad repeats within P*ctxAB* were tested to determine CT production in the presence and absence of HCO_3_^-^. As earlier reported (20), El Tor variant strain MCM168 and the Haitian variant IDH03595 (17) demonstrated heightened CT production compared to the prototype El Tor strain N16961, although the amount of CT produced varied among the strains. This enhanced CT production by the Haitian variant strain could also be attributed to the signature *ctxB7* allele, which results in a histidine to asparagine substitution (H20N) in the CT B signal sequence (17). Strikingly, MCM168, which has the minimal number of the TTTTGAT heptamer sequence within P*ctxAB*, produced the highest amount of CT *in vitro* even when HCO_3_^-^ was not included in the AKI medium. The classical strain H218 with 6 perfect heptad repeats was found to be severely attenuated in CT production under both of the conditions, although the mechanism underlying this particular phenotype remains elusive. O395 produced approximately twice the amount of CT as that produced by 569B in the absence of HCO_3_^-^, although it produced CT in equivalent amount in AKI medium supplemented with HCO_3_^-^. Given that the classical strains carry two copies of the *ctxAB* operon; one on each of the two chromosomes and the ToxT expression levels are usually higher in these strains than the prototype El Tor biotype strains (23), this may account for the increased CT production by O395 and 569B. We initially hypothesized that the differing amounts of CT produced by the tested strains (Figure 1 B) could be indirectly linked to the number of the heptad repeats, and the distal repeats could somehow contribute to H-NS repression.

For a more straightforward analysis, we then designed *ctx::lacZ* transcriptional fusions. These reporters ranged from +21 up to −257 relative to the TSS. The 5’ end of each fusion DNA spans a partial 3’ end of the upstream *zot* gene, the stop codon of which consists of the 5’ portion of the most distal TTTTGAT sequence. As the number of the heptad repeats varies among the two biotypes of *V. cholerae, ctx::lacZ* plasmids were selectively chosen for the P*ctxAB* activity analysis, with the minimum and maximum number of repeats for each biotype. Quantitative β-galactosidase activity assay in wild type El Tor strain N16961 demonstrated that the number of the repeats within P*ctxAB* negatively correlates with the expression levels from the fusion plasmids, indicating that the distal repeats in classical strains could possibly participate in H-NS binding along the A/T rich promoter sequence. This result was further validated by *in vitro* gel shift experiments where we could observe that a supershift appeared when purified H-NS from *V. cholerae* was added to DIG-labelled DNA with 5’ end point −243, therefore suggesting H-NS polymerization along the heptad repeats. Interestingly, this supershift was not prominently observed when DNA substrate shorter by a single heptad repeat was used as EMSA probe.

Our observation is further supported by previous studies (10, 11) which provided experimental evidences that H-NS utilizes the heptad repeats for binding to P*ctxAB*. However, this also raises a probability that full displacement of H-NS by ToxT may not be necessary for transcriptional activation by ToxT. In other words, even under virulence inducing condition, the degree of H-NS displacement from P*ctxAB* could depend on the number of the direct repeats within the regulatory region upstream of the *ctx* genes. Despite of our repeated attempts and those of collaborators, we could not make an *hns* mutant strain of *V. cholerae*. Therefore, further experiments are needed to understand how the number the heptad repeats upstream of the basal promoter element and within the *zot-ctxAB* intergenic region could correlate with H-NS repression and counter repression by ToxR and ToxT, which also recognize conserved motifs within P*ctxAB*.

In order to avoid *V. cholerae* specific factors that may have an interfering effect on P*ctxAB* activation, β-galactosidase reporter assays were performed with derivatives of *E. coli* K37. It was evident that in the presence of both H-NS repression and ToxT activation, P*ctxAB* activity inversely correlates with the number of TTTTGAT heptad repeats, at least partially. Interestingly; unlike our above-mentioned observation in *V. cholerae*, we found no change in *ctx::lacZ* expression among classical P*ctxAB* fragments with 5’ ends from −243 to - 257. Based on our results, we suspect that either H-NS has a specific binding region within P*ctxAB* that encompasses not all of the heptad repeats or not every repeat may equally contribute to H-NS binding. This was further supported by promoter activity experiments in WT *E. coli* with empty pMMB308 vector. In these strains, overall background levels of *ctx::lacZ* expression dropped sharply compared with *E. coli* strains expressing 6XHis-ToxT upon addition of IPTG as there is little *PctxAB* activity in the absence of ToxT. Interestingly, H-NS repression significantly elevated as the number of the heptad repeats in the fusion plasmids increased from 4 to 5, and no further reduction in promoter activity could be found even if the number of the repeats extended up to the maximum of 8. Expression of ToxT dramatically increased the overall transcriptional levels in *Δhns E. coli*, although ToxT-dependent P*ctxAB* activation did not differ among the fusion constructs with heptad repeats 4-8, and the shortest promoter still demonstrated the highest level of β-galactosidase activity. These results are in complete agreement with earlier observations (10, 11) that P*ctxAB* is repressed by H-NS, and ToxT counteracts by antirepression and stimulation of RNAP activity.

According to the current model of ToxT transcription activation (11), P*ctxAB*, is occupied by H-NS under virulence repressing condition (when ToxT initially is in an inactive state (19), and therefore *ctx* expression is transcriptionally silenced. Under virulence inducing conditions favourable for ToxT dependent transcriptional activation, ToxT displaces H-NS and binds to the two toxboxes within P*ctxAB*. The ToxT binding region comprises heptad repeats 6 and 7, which are most proximal to the transcription start site. ToxT then transcriptionally activates the *ctx* promoter, which results in the expression of toxin mRNA and subsequent CT production. However, as ToxT was not observed to bind to the distal repeats in footprinting experiments (Dittmer and Withey Unpublished data), and H-NS utilizes the heptad repeats to transcriptionally silence *ctxAB*, our recent understandings about H-NS and ToxT binding to P*ctxAB* and the studies presented here suggest that H-NS represses P*ctxAB* by occupying the distal heptad repeats in *V. cholerae*, and both H-NS and ToxT may occupy P*ctxAB* simultaneously, even under the virulence inducing condition when ToxT is in fully active form. Based on our data, we hypothesize that H-NS could be only displaced by ToxT from the proximal heptad repeats where H-NS and ToxT binding sites overlap, and therefore the number of the heptad repeats inversely correlate with the transcriptional activation of the *ctxAB* operon. Taken together, this study proposes a revision of the current model regarding H-NS repression and ToxT dependent transcriptional activation of P*ctxAB* (Figure 7). Further investigations could elucidate the exact role of the heptad repeats in controlling DNA-protein interactions between the A/T rich P*ctxAB* and other DNA binding proteins such as ToxR and RNAP in presence of H-NS and ToxT. The outcome will be significant because of new information about transcriptional regulation of virulence gene promoters in *V. cholerae* and enteric pathogens.

**Figure 7.**
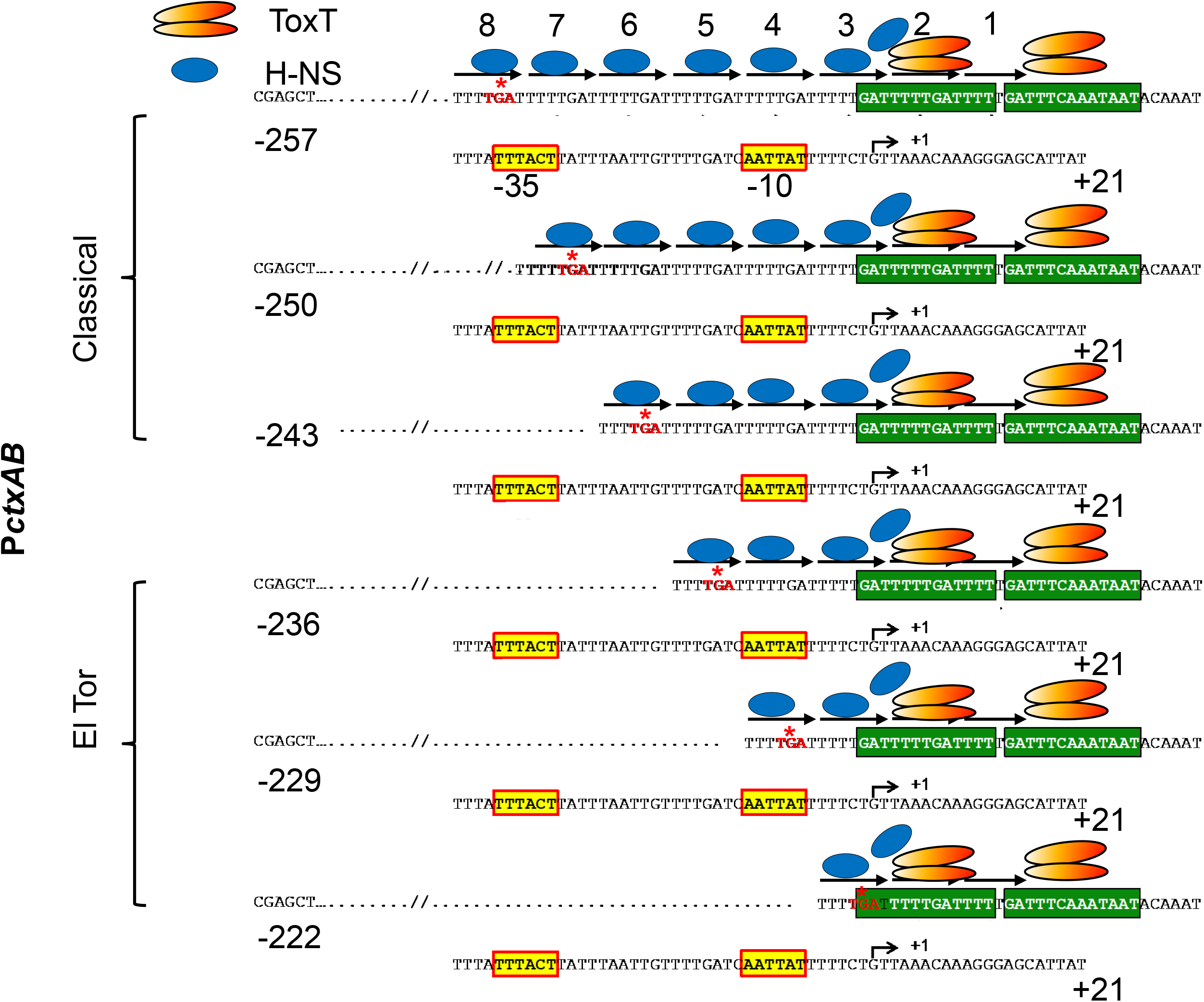
Proposed revised model of P*ctxAB* transcriptional regulation. Model for transcriptional activation El Tor and classical *PctxAB* regions (starting from +21 with 5’ end up to −257) is summarized. The ToxT binding sites are indicated by green boxes. The bend arrow represents the transcription start site. Putative −10 and −35 core promoter elements are highlighted within yellow boxes. The red asterisk indicates TGA stop codon of the *zot* gene. TTTTGAT heptad repeats are marked with black arrows. In the absence of ToxT dependent activation, H-NS utilizes the heptad repeats and polymerize along the A/T rich sequences within P*ctxAB*. Under virulence inducing conditions, ToxT binds to the toxboxes and displace H-NS monomers from toxbox 1. Any other H-NS oligomers which are occupied over the distal heptad repeats, however remain bound to the P*ctxAB* and interfere with the promoter activation of the cholera toxin operon.

## ACKNOWLEDGEMENTS

Part of this work has been presented at the “International Symposium on Systems, Synthetic & Chemical Biology (SSC 2017)” We thank Dr. Rupak K. Bhadra (CSIR-Indian Institute of Chemical Biology) for providing us the *V. cholerae* H218 strain.

This research was supported in part by the Indian Council of Medical Research (ICMR); **the Japan Agency for Medical Research and Development (AMED) [grant number JP21wm0125004 (to SM)];** and the National Institute of Infectious Diseases (NIID), Japan. AG received JC Bose Chair Professorship of The National Academy of Sciences, India (no. NAS/2057/3/2015-16, URL: http://www.nasi.org.in/). AN and PM received Indian Council of Medical Research (ICMR) fellowships (No. ICMR-SRF No. 80/890/2014-ECD and No. 80/748/2012-ECD-I), P.S. and R.N.S. acknowledge Council of Scientific and Industrial Research (CSIR) fellowship [No. 09/482(0060)/2014-EMR-I and ICMR fellowship (No. 3/1/3/JRF-2015/HRD-LS/88/40189/82), respectively.

## AUTHOR CONTRIBUTIONS

A.N. designed the research and A.K.M. supervised the study. A.N., J.H.W., and A.K.M. conceptualized the study. A.N., J.H.W., and A.K.M designed the experiments. A.N. carried out the experiments. P.M. participated in molecular cloning. R.N.S. participated in the CT ELISA and reporter assay experiments. P.S. participated in EMSA experiments. A.N. and J.H.W analyzed the data. J.H.W., SM, and S.D. contributed new reagents/analytic tools. AN. wrote the paper and J.H.W., A.G. and A.K.M contributed in writing. All authors have gone the manuscript and approved for submitting to the journal.

---

**Figure S1. Purification of the *V. cholerae* H-NS protein by chitin affinity column.**

C-terminally intein tagged H-NS protein was purified using the IMPACT kit as described under “materials and methods”. Following thiol group mediated cleavage by the addition of DTT, eluted fractions were collected and analyzed by 15% SDS-PAGE. Positions of the marker proteins (PageRuler^™^ Prestained Protein Ladder, Catalogue No. 26617) of known molecular weights (kDa) are indicated.

