## Supplementary figures and images for "Elucidating the correlation between the number of TTTTGAT heptamer repeats and cholera toxin promoter activity in *Vibrio cholerae* O1 pandemic strains"

### Supplemental File 1

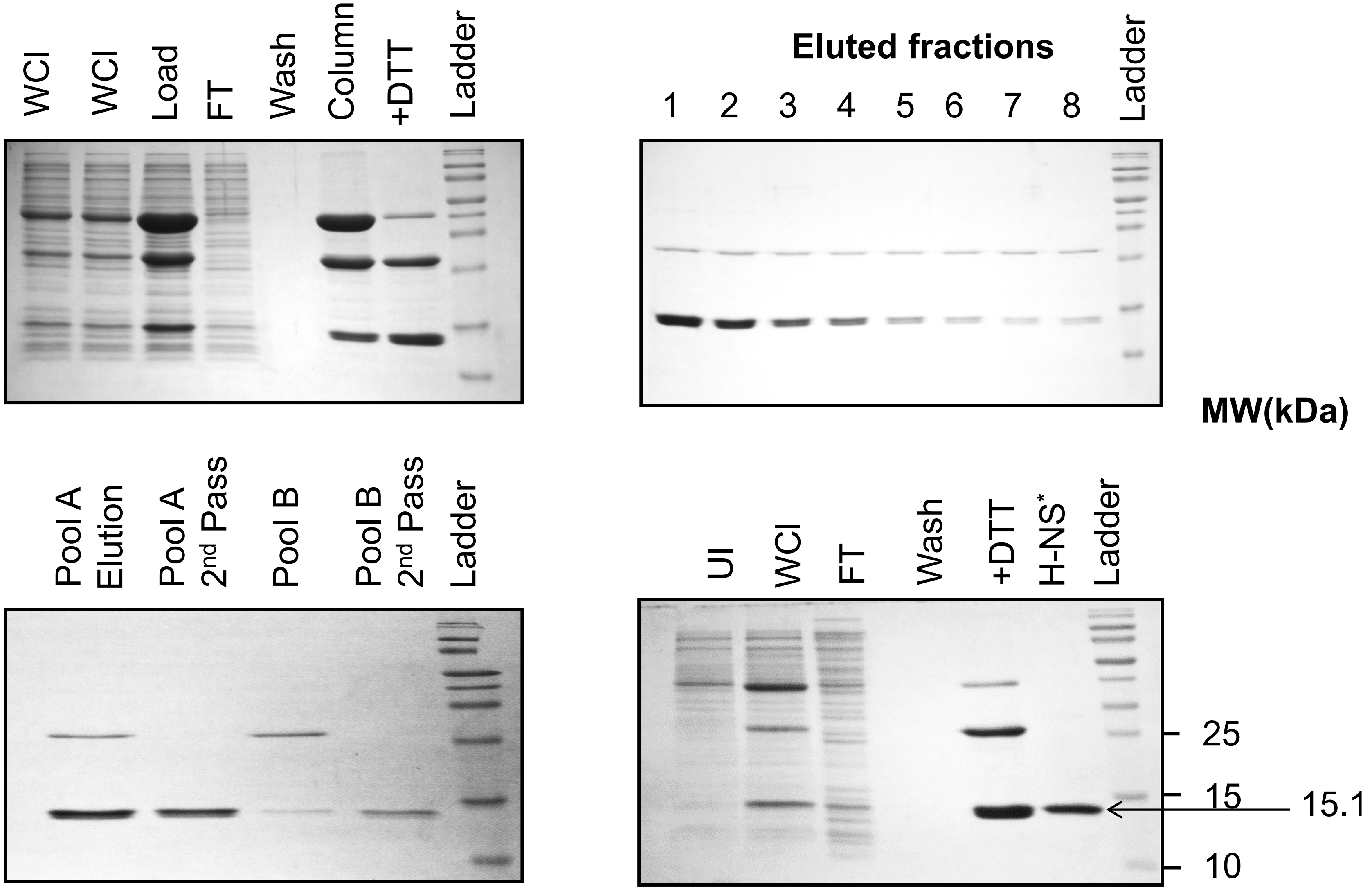


* After dialysis and addition of glycerol to final 10%

Fig.S1
